# Glucagon-like peptide-1 receptor agonist, exendin-4, reduces reinstatement of heroin seeking behavior in rats

**DOI:** 10.1101/730408

**Authors:** Joaquin E. Douton, Corinne Augusto, Brooke A Stultzfus, Nurgul Carkaci-Salli, Kent E. Vrana, Patricia S. Grigson

## Abstract

**Background:** Studies have shown that ‘satiety’ agents such as exendin-4 (a glucagon-like peptide-1 analog) reduce responding for addictive drugs (e.g., cocaine, nicotine, alcohol). In this study we tested the effect of exendin-4 on cue-induced and drug-induced reinstatement of heroin seeking behavior in rats.

**Methods:** This study consisted of three phases: In Phase 1, 55 male Sprague-Dawley rats had 15 daily pairings of saccharin with heroin self-administration. In Phase 2, rats experienced a 16-day home cage abstinence period and daily treatment with vehicle or exendin-4. On day 17, an extinction/reinstatement test was performed to assess drug seeking. In Phase 3, rats experienced 9 days of extinction followed by a reinstatement only test. Finally, expression of mRNA for various receptors in the nucleus accumbens shell (NAcS) was measured using RTqPCR.

**Results:** In Phase 1, rats that avoided intake of the heroin-paired saccharin cue exhibited shorter latency to obtain the first infusion. In Phase 2, treatment with exendin-4 decreased cue-induced, but not drug-induced heroin seeking. In Phase 3, saccharin avoiders previously treated with exendin-4 increased acceptance of saccharin, and 1-hour pretreatment with Exendin-4 abolished drug-induced heroin seeking. Finally, exendin-4 treatment increased expression of mRNA for the Orexin 1 receptor (OX1) in the NAcS, but did not affect expression of dopamine D2 receptors, GLP-1 receptors, or leptin receptors in this same structure.

**Conclusion:** Exendin-4 reduced cue- and drug-induced heroin seeking and increased acceptance of the drug-associated saccharin cue. These changes in behavior were accompanied by an increase in the expression of the OX1 receptor in the NAcS.

## INTRODUCTION

Drug addiction is a chronic disease that is difficult to treat due to its relapsing nature [1]. Overdose-related deaths tripled from 1999 to 2014 and, in 2017, about 70% of drug overdose deaths involved opioid use [2]. Currently, misuse of prescription pain relievers, heroin, and synthetic opioids causes the death of more than 130 people a day in United States alone, a tragedy of epidemic proportion and a major concern for the Centers for Disease Control and Prevention (CDC) [3]. It is, then, imperative to find effective treatments for drug addiction, more specifically, to mitigate the craving and withdrawal that precipitates relapse and, thereby, vulnerability to opioid overdose [4, 5].

Addiction is recognized as the pathological usurpation of neural systems associated with reward-related learning [6-10]. In this context, drug becomes the prominent goal, and in turn, natural rewards are devalued. This shift in motivation can be observed in individuals with substance use disorder (SUD) who fail to provide care to their children [11], show loss of productivity in the workplace [12], and decreased sensitivity to monetary rewards [13, 14]. Reward devaluation also has been observed in animal models, as cocaine-exposed female rats show greater preference for drug-associated stimuli than for their pups [15] and hungry and thirsty rats avoid intake of a palatable solution when it predicts the availability of a drug of abuse [16-20]. This insight led to the first animal model to study drug-induced devaluation of natural rewards [21, 22]. In this model, rats that most greatly avoid intake of a saccharin cue that predicts the availability of drug, show greater aversive reactions when the taste cue is intra-orally infused, show greater drug-seeking, drug-taking, willingness to work for the drug, and greater susceptibility to cue- and drug-induced relapse [18, 23-26]. Moreover, avoidance of the taste cue is associated with blunted levels of dopamine [27], high levels of corticosterone [28], and loss of body weight [29]. Collectively, these findings suggest that, by serving as a predictor of imminent drug access, the taste cue elicits the onset of a conditioned aversive state involving craving and/or withdrawal that can be corrected only by the taking of drug [21].

We have postulated that the metabolic state associated with craving a drug of abuse is similar to the state observed in animals seeking food when starved, water when thirsty, and salt when sodium-deficient. When such a state is reached, there is one goal and there is no substitute [21]. Such motivated behaviors are guided by the internal state of the individual and persist until the goal is obtained and an optimal homeostatic state is attained [30]. In this way, behavior healthfully switches from one motivated state to another depending on the needs of the system at any particular time. Drugs of abuse, on the contrary, engage one motivated behavior (i.e., drug seeking/taking) to the exclusion of the others [31].

That said, scientists have shown that hormones that modulate hunger and satiety can modulate responding for drugs of abuse [32-34]. The most studied so far has been Glucagon-like peptide-1 (GLP-1), an incretin hormone [35] essential for regulation of food intake in both animals and humans. GLP-1 also is a neurohormone released by neurons located in the nucleus tractus solitarius (NTS) that project to different brain regions, including key structures in the reward pathway such as the ventral tegmental area (VTA) and the nucleus accumbens (NAc) [36-39]. Exendin-4 (Ex-4) is a natural GLP-1R agonist and, due to its incretin and appetite suppressant action, has been used to treat type 2 diabetes mellitus and obesity [40-43]. Exogenous treatment with Ex-4 also has been observed to reduce conditioned place preference (CPP) and accumbens dopamine release by nicotine, cocaine and amphetamine; and to reduce cocaine self-administration and seeking in rats [44-49] and relapse-like drinking in mice [50]. Thus far, only one study investigated the effect of GLP-1R agonists on opioid addiction [51], finding Ex-4 not to be effective in reducing abuse-related effects of the opioid remifentanil in mice.

Not expecting opioids to be an exception, the present study used the model of reward devaluation to reexamine the effects of chronic treatment with Ex-4 on the response to a natural reward, cue-induced heroin seeking, and heroin-induced reinstatement of heroin seeking behavior (i.e. relapse) in rats. In addition, we analyzed mRNA expression in the NAc shell (NAcS), a crucial structure in reward modulation, to assess long-lasting changes in receptors associated with homeostatic regulation and reward. We hypothesize that chronic treatment with Ex-4 will reduce cue- and drug-induced heroin-seeking behavior following a period of abstinence, and will facilitate recovery of responding for the natural reward.

## METHODS AND MATERIALS

Fifty-five male Sprague-Dawley rats were implanted with jugular catheters and were trained and tested in self-administration chambers where they had access to three spouts. For more information about subjects, surgeries, solutions and apparatus see Supplemental Information.

### Phase 1: Acquisition of heroin self-administration behavior

#### Taste-drug pairings

Two hours into the light cycle, rats were placed in the self-administration chambers where they had 5-minute access to 0.15% saccharin via the leftmost spout. Thereafter, the house light was turned off and the middle and right empty spouts advanced. Licking on the inactive spout (middle spout) did not have any consequence. The active spout (rightmost spout) was signaled by a cue light located above and completion of a fixed ratio of 10 (FR=10) licks on this spout led to a 6-second iv infusion of either saline (n=15) or 0.3 mg/infusion heroin (n=40) for a period of 6 hours. Each infusion was followed by a 20-second time-out period during which the cue light turned off, the house light turned on, the spouts retracted and the sound of a tone signaled drug unavailability. Rats were trained using this protocol 5 days a week for a total of 15 trials (Figure1). Due to lack of catheter patency, seven animals in the saccharin-heroin group were excluded from the experiment after Phase 1.

**Figure 1.**
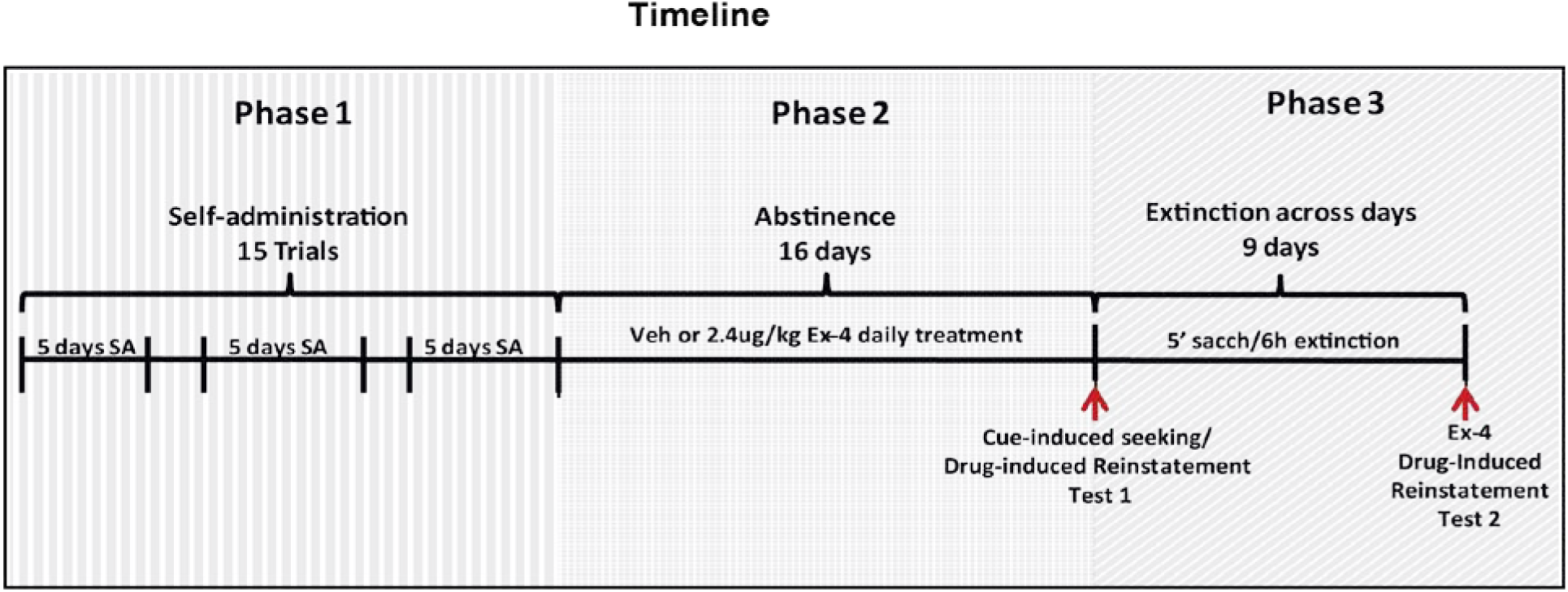
Timeline of the study. The study consisted of 3 phases. During Phase 1, rats had 5 min access to a saccharin cue followed by the opportunity to self-administered either heroin or saline 5 days a week for 15 trials. Phase 2 consisted of a 16-day period of home cage abstinence followed by Cue-induced seeking/Drug-induced Reinstatement Test 1. During this period rats were treated with a daily ip injection of either Veh (saline) or 2.4µg/kg Ex-4. During phase 3 rats were subjected to a daily paring of saccharin with a 6-h extinction period for 9 days. During Drug-induced Reinstatement Test 2, the rats were given 5 min access to the saccharin cue, followed by three hours of extinction testing. The small and large suppressor rats were injected ip with Veh or Ex-4 at the end of h 2, the iv heroin prime injection was infused at the end of h 3, and reinstatement of heroin seeking behavior was examined for 1 h thereafter.

#### Behavioral stratification

In this paradigm, some rats avoid the drug-paired saccharin cue and greater avoidance is associated with greater drug-seeking and taking [18, 52]. Consequently, we used terminal saccharin intake (i.e. mean saccharin licks in trials 14 and 15) to separate the heroin self-administration group into large and small suppressors. Large suppressors (LS) emitted, on average, <200 licks/5min (n=19), while small suppressors (SS) emitted >200 licks/5min (n=21) during trails 14 and 15. Saccharin-saline (Sac-sal) controls were not stratified.

### Phase 2: Abstinence, treatment and drug-seeking tests

#### Abstinence

A 16-day abstinence period followed immediately after the last taste-drug pairing. During this period, rats remained in their home cage with ad-lib access to water and food and were injected intraperitoneally (ip) once daily with either vehicle (Veh) or 2.4µg/kg of Ex-4 one hour into the light cycle.

#### Cue-induced seeking and drug-induced reinstatement test 1

During this test, rats received their respective Vehicle or Ex-4 injection one hour prior to placement in the chamber. They then had 5 minutes access to saccharin followed by a 5-hour extinction test during which all cues associated with the drug (cue light, tone, etc.) were presented, but contacts on the active spout did not deliver infusion of drug. Immediately after the fifth hour, a non-contingent iv heroin infusion was automatically delivered by the computer and subsequent contacts with the spouts were recorded to assess drug-induced reinstatement of heroin seeking behavior. Due to lack of catheter patency, five additional rats in the saccharin-heroin group were excluded of the experiment after Phase 2. In addition, one rat in the Sac-sal group developed an infection and had to be euthanized.

### Phase 3: Extinction across days and drug-induced reinstatement test 2

In Phase 2, our standard protocol led to a 6-hour delay between Ex-4 treatment and the drug-prime. Since, Ex-4 has a half-life of around 150 minutes [53] by the time the drug prime was delivered, the levels of Ex-4 in the system are expected to be low. In order to more effectively assess the impact of Ex-4 on drug-induced reinstatement, rats were subjected to a protocol of extinction across days to reduce cue-induce seeking, and Ex-4 was administered 1 hour prior to drug-induced reinstatement test 2.

#### Extinction across days

Two hours into the light cycle rats were placed in the chamber and given 5 minutes access to saccharin, followed by a 6-hour extinction test in which contacts with the active spout were without consequence. During this 9-day period no Ex-4 or vehicle was administered.

#### Drug-induced reinstatement test 2

On day 10, and two hours into the light phase, the rats had 5 minutes access to saccharin. Because cue-induced seeking was not fully extinguished across the 9-day period, rats were exposed to 3 hours of extinction within session on Day 10. At the beginning of hour 2, rats were injected ip with either saline or Ex-4. One hour thereafter, rats received a non-contingent iv infusion of heroin and reinstatement of heroin seeking was examined across the next hour. The Sac-sal group was not subjected to this final test.

#### Sacrifice, dissection and molecular analysis

One hour after the last test, all rats were sacrificed by live decapitation, the skull was removed and the brain dissected. The harvested right NAcS was flash frozen and stored at −80 °C until mRNA was extracted and analyzed by real time quantitative polymerase chain reaction (RT-qPCR). For more details see Supplemental Information.

## RESULTS

### Phase 1: Acquisition of heroin self-administration behavior

#### Saccharin intake

As shown in Figure 2A, the number of saccharin licks/5min increased across trials in both Sac-sal and SS, but not LS. Support for this conclusion was provided by post-hoc assessment of a significant 3 × 15 mixed factorial ANOVA varying group (Sac-sal, SS and LS) and trials (1-15) (F_28,728_=10.52, p<0.0001) showing a significant reduction in intake of the saccharin cue on trials 7 to 15 for the LS vs. both the Sac-sal and the SS (ps<0.05).

**Figure 2.**
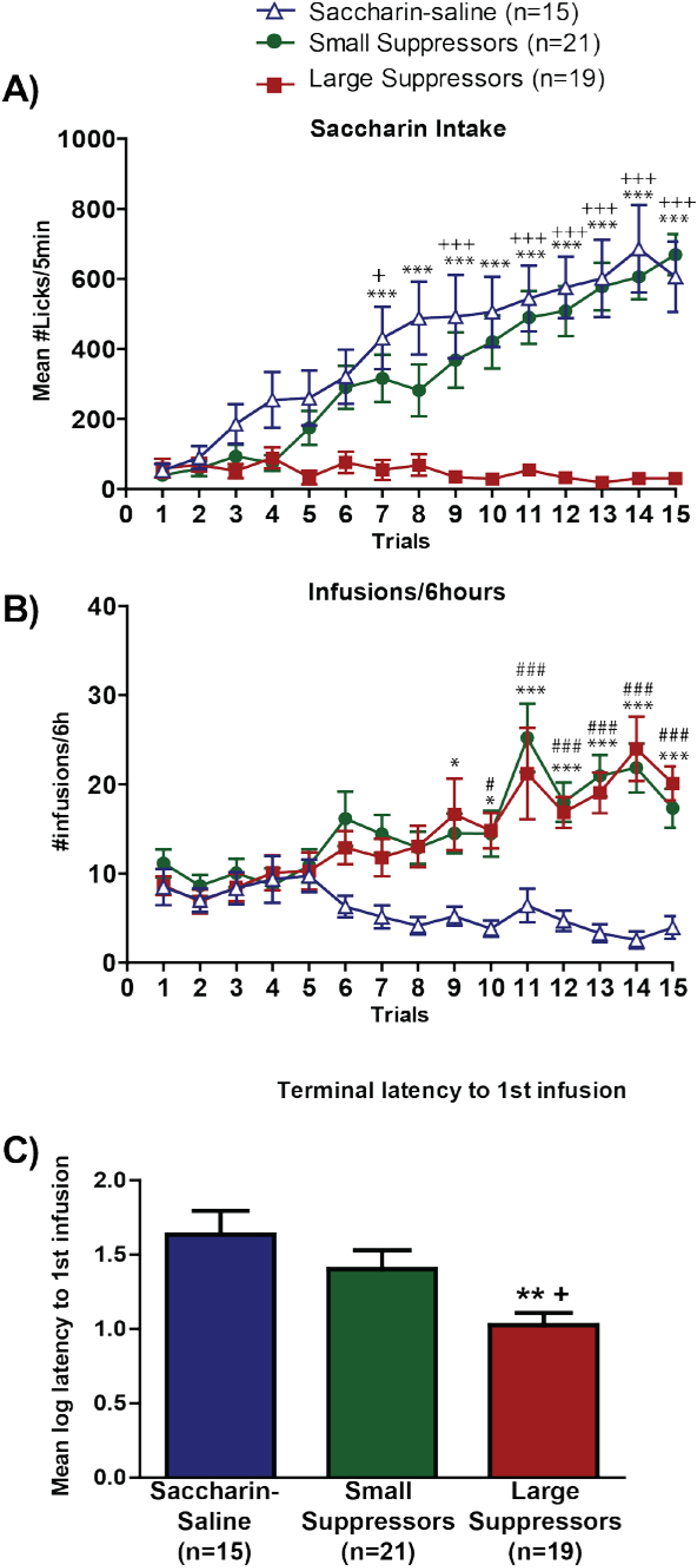
Acquisition of drug-taking behavior. **A)** Mean± SEM number of licks/5min of 0.15% saccharin across trials 1 – 15 for rats in the saccharin-saline, small suppressor and large suppressor group. **B)** Mean± SEM number of heroin infusions/6h across trials 1 – 15 for rats in the saccharin-saline, small, and large suppressor groups. **C)** Mean± SEM terminal log latency to 1^st^ infusion for saccharin-saline, small, and large suppressor groups. (saccharin-saline controls: n=15, small suppressors: n=21, large suppressors: n=19). ^+^Significant difference between large suppressors and small suppressors; ^*^Significant difference between large suppressors and saccharin-saline controls. # Significant difference between small suppressors and saccharin-saline controls. For any given symbol: _*_<0.05; _**_<0.01; _***_<0.001.

#### Heroin self-administration

There were no significant differences in the mean number of infusions/6 hours across the 15 trials of self-administration behavior between SS and LS, and both groups self-administered more infusions than the Sac-sal (Figure 2B). This conclusion was supported by a 3 × 15 mixed factorial ANOVA showing a significant Group x Trial interaction (F_28,546_=3.18, p<0.0001). Post-hoc tests revealed a greater number of heroin infusions by both the SS and LS compared to number of saline infusions by Sac-sal, beginning on trial 9 for the LS and on trial 10 for the SS (ps<0.05). In addition, LS evidenced a shorter terminal latency, averaged across trials 14 and 15, to obtain the first infusion (Figure 2C) compared with SS and Sac-sal. This was confirmed by post-hoc analysis of a One-Way ANOVA (F_2,51_=5.78; p=0.005, ps<0.05).

### Phase 2: Abstinence, treatment and drug-seeking tests

#### Saccharin intake after abstinence

Ex-4 treatment during abstinence did not affect body weight (see Supplemental Figure 1) or saccharin intake at test (Figure 3A). Number of licks/5min during test 1 was analyzed using a 3 × 2 ANOVA varying group (Sac-sal, SS and LS) and treatment (Veh vs. Ex-4). The results revealed a significant main effect of group (F_2,40_=29.29, p<0.0001), with LS group emitting fewer saccharin licks/5min than SS or Sac-sal (p<0.05). There was, however, no significant main effect of treatment (F_1,40_=1.77, p=0.28) or group x treatment interaction (F<1.0).

**Figure 3.**
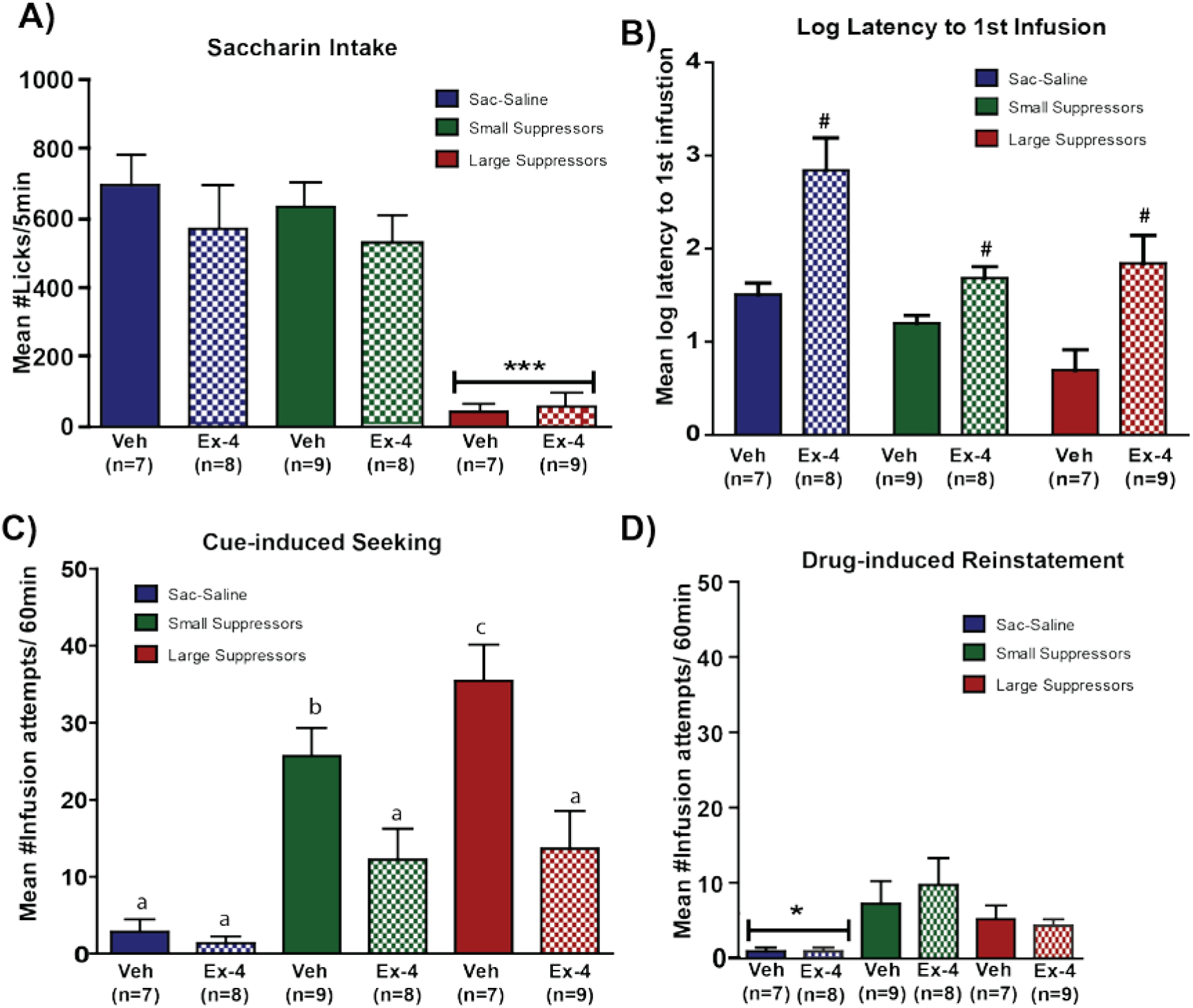
Effect of Ex-4 treatment on cue- and drug-induced reinstatement. **A)** Saccharin intake on test day. Mean ± SEM number of licks/5min of 0.15% saccharin for rats in the saccharin-saline controls, small suppressors and large suppressors groups treated with Veh or Ex-4 on cue-induced seeking and drug-induced reinstatement test 1. **B)** Latency to 1^st^ infusion attempt. Mean± SEM log latency to 1^st^ infusion attempt for saccharin-saline, small and large suppressor groups treated with Veh or Ex-4. **C)** Cue-induced seeking. Mean± SEM number of infusion attempts/60min for saccharin-saline, small and large suppressors groups treated with Veh or Ex-4 during the first hour of seeking. **D)** Drug-induced reinstatement test 1. Mean± SEM number of infusion attempts during the 60 minutes after the drug prime for saccharin-saline, small and large suppressors groups treated with Veh or Ex-4 six hours prior to the drug prime. *Significant difference between groups.*p<0.05, ***p<0.001; #Significant difference between veh and Ex-4 treatment. Different letters indicate significant differences

#### Cue-induced Seeking

Figure 3B shows that treatment with Ex-4 increased the mean log latency to the first infusion attempt. In support, the 3 × 2 ANOVA varying group and treatment found a significant main effect of treatment (F_1,41_=29.83, p<0.001). The main effect of group also attained statistical significance (F_2,41_=9.116, p=0.001), while the group x treatment interaction did not (F_2,41_=2.12,p=0.132). Post-hoc analysis of group (ps<0.005) confirmed a longer latency to the first infusion attempt by Sac-sal vs. SS and LS (p < 0.05).

Exendin-4 significantly reduced infusion attempts during the 1^st^ hour of seeking after abstinence (i.e. during cue-induced reinstatement) in both LS and SS (Figure 3C). Support for this conclusion was provided by a significant group x treatment interaction (F_2,40_=8.53, p<0.001) and confirmed by a post-hoc analysis (ps<0.05).

#### Drug-induced reinstatement test 1

Treatment with Ex-4 six hours earlier had no effect in the number of infusions attempted during the hour following the heroin prime (Figure 3D). While Sac-sal rats treated with Veh or Ex-4 did not attempt any infusions, both vehicle-treated SS and LS showed a higher number of infusion attempts than Sac-sal. Thus, a 3 × 2 ANOVA showed a significant main effect of group (F_2,38_=9.48, p=0.0005) and a post-hoc analysis showed significantly greater infusion attempts by both SS and LS groups vs. Sac-sal (p<0.05). Neither the main effect of treatment (F<1.0) nor the group x treatment interaction (F<1.0) was significant.

### Phase3: Extinction across days and drug-induced reinstatement test 2

#### Saccharin intake across extinction days

A history of treatment with Ex-4 significantly increased the number of saccharin licks/5min across 9 days of extinction in the LS (Figure 4A). In support, a 2 × 3 ANOVA revealed a significant group x treatment interaction (F_2,48_=3.97, p=0.02). Post-hoc tests (ps<0.005) confirmed a significant increase in saccharin intake by the Ex-4 vs. Veh-in the LS and Sac-sal groups.

**Figure 4.**
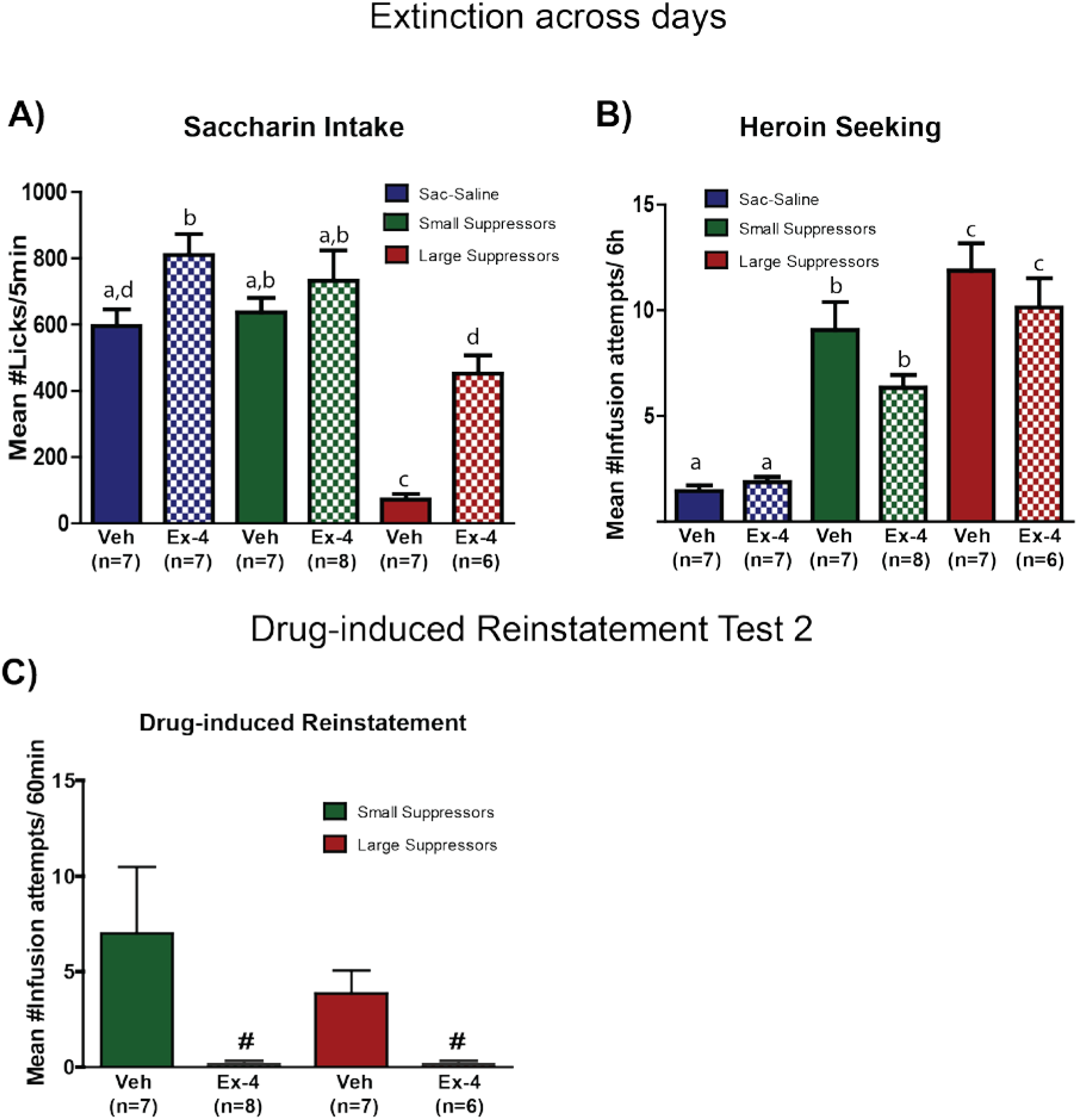
Extinction across days and drug-induced reinstatement test 2. **A)** Saccharin intake during extinction. Mean± SEM number of licks/5min of 0.15% saccharin averaged across the 9 days of extinction for rats in the saccharin-saline, small suppressors and large suppressors groups with history of treatment with Veh or Ex-4. **B)** Heroin seeking during extinction. Mean ±SEM number of infusion attempts/6h averaged across 9 days of extinction testing for saccharin-saline, small and large suppressors with a history of treatment with Veh or Ex-4. **C)** Drug-induced reinstatement test 2. Mean± SEM number of infusion attempts/60min for small and large suppressors treated with Veh or Ex-4 1 hour prior to the drug prime. #Significant difference between veh and Ex-4 treatment. Different letters indicate significant differences.

#### Drug seeking across extinction days

History of treatment with Ex-4 did not have an effect on number of infusion attempts/6h during the 9-day extinction period (Figure 4B). This conclusion is confirmed by a 3 × 2 ANOVA showing no significant main effect of treatment (F_1.44_=2.39, p=0.128) or group x treatment interaction (F_2,44_=1.05, p=0.36). A significant main effect of group (F_2,44_=37.15, p<0.0001) and post-hoc analysis (ps<0.005) showed a significant difference between the three group..

#### Drug-induced Reinstatement Test 2

Figure 4C shows that acute-pretreatment with Ex-4 completely abolished heroin seeking during the heroin-induced reinstatement test 2. Vehicle-treated SS and LS showed an increased number of infusion attempts, while groups pre-treated with Ex-4 did not attempt any infusions at all. This conclusion was supported by a 2 × 2 ANOVA showing a significant main effect of treatment (F_1,21_=7.15;p=0.014). Neither the main effect of group (F<1.0), nor the group x treatment interaction (F<1.0), were significant.

#### Molecular Analysis

To further explore the effects of Ex-4, mRNA expression of different genes associated with reward and feeding behaviors were examined in the NAcS. Figure 5 shows the relative expression of 4 such genes in rats that self-administered heroin and were treated with Veh or Ex-4. No significant difference in fold change expression was observed in GLP-1R (Figure 5A), D2R (Figure 5B), or leptin receptors (Figure 5C). However, the expression of Orexin receptor 1 (OX1) (Figure 5D) was significantly higher in heroin self-administering rats with a history of Ex-4 treatment compared to vehicle-treated controls (t=4.56; p=0.001).

**Figure 5.**
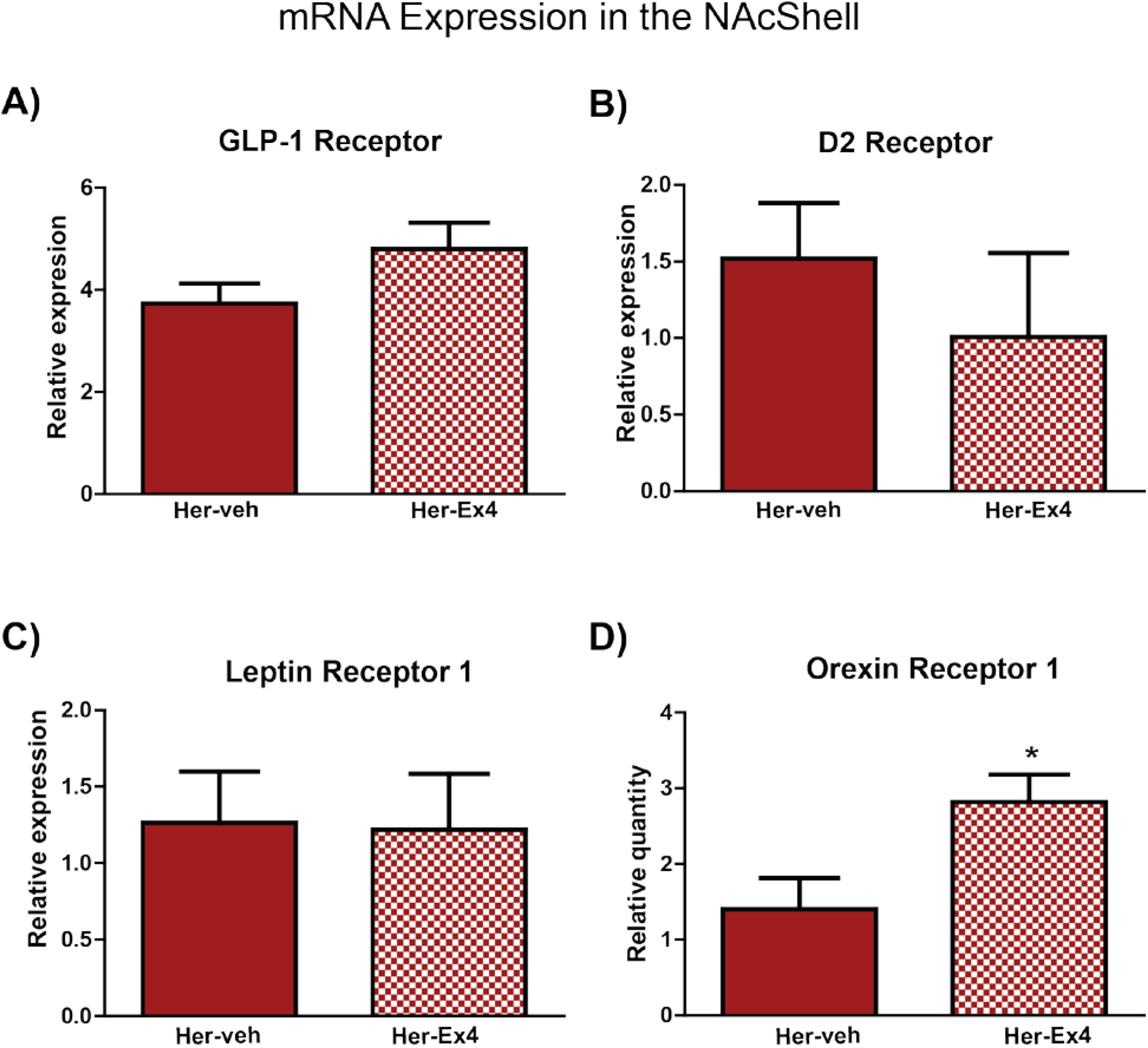
mRNA expression. Relative mRNA expression of GLP-1 receptor **(A)**, dopamine D2 receptor **(B)** leptin receptor 1 **(C)** and orexin receptor 1 **(D)** in rats with a history of heroin self-administration and treatment with Veh or Ex-4 throughout the abstinence period and test. The n indicates the number of biological replicates. *p<0.05

## DISCUSSION

This study examined the effects of the Ex-4 on heroin addiction-like behaviors in rats. Specifically, we showed that chronic, systemic treatment with Ex-4 during abstinence significantly attenuated heroin seeking when the animals were re-exposed to the heroin-associated cues without affecting body weight. Moreover, Ex-4 abolished drug-induced reinstatement of heroin seeking behavior when administered 1 hour, but not 6 hours, prior to the drug challenge. Furthermore, a history of treatment with Ex-4 increased acceptance of the heroin-paired saccharin cue, particularly in the most vulnerable LS population. Finally, reduced reinstatement of heroin seeking by Ex-4 treated rats was accompanied by an increase in the expression of the OX1 mRNA in the NAcS.

### Individual differences

In Phase 1 of the present study we did not observe differences in heroin taking between SS and LS. As mentioned, it is well established that rats will avoid intake of a taste cue when paired with drugs of abuse [17, 18, 20, 26, 54, 55], and greater avoidance can predict greater susceptibility to drug addiction [18, 24]. Although this is true for several drugs, the differences in drug-taking between SS and LS using heroin are, at times, not as robust as those observed with cocaine, for example [18, 25, 52]. That said, in this model, the taste cue is thought to elicit a conditioned withdrawal state of ‘need’ [21]. Thus, a better measure of this ‘need’ state may be the latency to obtain the first infusion of drug. In accordance, while the SS and LS rats self-administered the same amount of heroin, LS took less time to obtain the first infusion, consistent with a greater state of ‘need’ in this subpopulation.

### GLP-1R agonist effects on drug seeking

In phase 2 of this study, we showed that chronic treatment with 2.4 µg/kg of Ex-4 did not affect body weight. Moreover, it reduced cue-induced but not drug-induced seeking of heroin. As mentioned, the injection of Ex-4 was delivered more than twice the half-life of the drug prior to the heroin prime. Although the expectation of the treatment protocol was to achieve long-lasting effects, the protective properties of this dose of Ex-4 clearly did not carry over across days. For this reason, in Phase 3, the effects of Ex-4 were tested directly on drug-induced reinstatement by reducing the pre-treatment time to one hour. Using this protocol, we showed for the first time that Ex-4 completely abolished heroin-induced reinstatement of heroin-seeking behavior (i.e., ‘relapse’) in both SS and LS.

### Effects of Ex-4 on drug-paired saccharin intake

As stated in the Introduction, part of our hypothesis was that the reduction in heroin seeking would be accompanied by an increase responding for the natural reward. This hypothesis was partially correct, since a history of chronic treatment with Ex-4 did increase acceptance of the saccharin solution in group LS, but only during extinction training when the saccharin cue was paired with the absence of heroin self-administration. In addition, while this Ex-4 dose can reduce responding for sucrose [56], those effects were not observed using saccharin in this experiment, as evidenced in the Sac-sal and SS groups. Moreover, high responses to saccharin suggest that Ex-4 has a specific effect on drug-seeking that is not due to a general reduction in locomotor activity. Overall, increased responding for saccharin in the LS is a promising finding, as natural rewards are our best natural defense against addiction. Future studies will need to determine whether extended treatment with Ex-4 restores the perceived palatability of the saccharin per se, or increased intake by reducing the onset of cue-induced craving and/or withdrawal.

### Molecular data

In the molecular analysis, we showed that treatment with Ex-4 increased mRNA expression for OX1 in the NAcS, but did not change the expression for GLP-1R, D2R or LepR1. The integrative nature of NAc, receiving inputs from different parts of the brain, including regions that control reward and feeding-motivated behaviors [57, 58], makes it a candidate area to look for long-term changes produced by Ex-4 treatment. Indeed, GLP-1 producing neurons in the NTS project to the NAc [59]. Moreover, systemic administration of Ex-4 reaches the NAc, and infusion of a GLP-1R agonist directly into the NAcS dose-dependently reduces cocaine seeking [60] without affecting food intake [61]. It was therefore expected that prolonged, chronic treatment with Ex-4 may affect the expression of GLP-1R in the NAcS. This, however, was not observed by qPCR analysis.

Another important source of input to the NAc are the dopaminergic projections from the VTA. GLP-1R are found in this area [36] and intra-VTA injections of Ex-4 have been shown to reduce drug-primed reinstatement of cocaine-seeking behavior. In addition, systemic injections of Ex-4 were found to attenuate amphetamine-, cocaine-, and nicotine-induced accumbal dopamine release [45, 62], showing that activation of GLP-1R can modulate drug-associated dopaminergic release in the NAc. Moreover, the protective effects of systemic Ex-4 on cocaine-seeking are blocked by infusion of a GLP-1R antagonist directly into the VTA [49]. Interestingly, we did not find changes in the expression of D2 receptors in the NAcS associated with Ex-4 treatment.

Finally, we focused on the expression of receptors associated with the homeostasis-related hormones leptin and orexin. Leptin is an adipose-derived peptide hormone that can attenuate drug-related behaviors such as cocaine-induced CPP, cocaine seeking and stress-induced reinstatement of heroin seeking [34, 63, 64]. However, in this experiment, Ex-4 did not have an effect on LepR1 mRNA expression in the NAcS. On the other hand, we did see a marked increase in the expression of OX1 in the NAcS. This receptor is exclusively activated by Orexin A [65], which is produced mainly by neurons within the lateral hypothalamus [65, 66], and has an important role in arousal and feeding motivated behavior. Orexins increase food intake [66, 67], but its modulating role also has been extended to non-feeding reward-related processes such as addiction [68, 69]. Our results suggest that treatment with Ex-4 might reduce the release of orexin in the NAcS, leading to an increase in the expression of the OX1. This is consistent with previous research showing that GLP-1Rs in the lateral hypothalamus are critical for the control of ingestive behaviors, body weight, and food reinforcement in rats [70]. In addition, blocking the effects of OX1 with systemic administration of OX1 antagonist SB-334867 dose-dependently blocked reinstatement of seeking for cocaine [71], alcohol [72, 73], nicotine [74] and remifentanil [75]. Future studies will need to parse the roles of the GLP-1R and the OX1 in the eventual acceptance of the natural reward and in the reduction in both cue- and drug-induced reinstatement of heroin-seeking behavior.

### Inconsistent findings

The effects of Ex-4 on opioid addiction have been addressed in only one study so far [51]. Contrary to our findings, Bornebusch et al. did not find Ex-4 to be effective in protecting mice from remifentanil seeking and taking. Besides using different subjects (mice vs. rats) and a different drug (remifentanil vs. heroin), they performed their studies in the dark, while our study was conducted during the light phase of the cycle. It has been shown that heroin access during the light phase leads to a full reversal of sleep/wake cycle [76]. If the effects of Ex-4 are dependent on the circadian activity of orexin neurons, the time of treatment or testing might be crucial for the effectiveness of the drug [77, 78]. Bornebusch et al. did not observe effects of Ex-4 on self-administration using FR1, FR3 or FR5, however they did see a trend for central GLP-1 KO mice to show higher responding on a progressive ratio paradigm. In addition, they did not see Ex-4 effects on CPP when administered during the conditioning phase. This is consistent with the hypothesis that Ex-4 acts via the orexin system to reduce opioid seeking. The orexin system is engaged in situations of high motivational relevance [79], hence it is expected to be involved during expression rather than acquisition of CPP [80, 81], or in self-administration when the task requires more effort (FR>5 or PR) [81, 82]. If this interpretation is correct, then treatment with Ex-4 is effective in our paradigm that requires higher motivation (FR10), but not when using FR<5. Further experiments need to be performed to elucidate the true cause of the discrepancy and to address the extent to which the effects of Ex-4 are dependent on the orexin system.

## CONCLUSION

Our results demonstrate that treatment with the GLP-1R agonist, Ex-4, attenuates cue-induced heroin seeking and drug-induced reinstatement of heroin seeking in rats. The molecular results are consistent with the notion that drugs of abuse not only highjack the reward system, but also generate a homeostatic dysregulation that is associated with a compelling state of physiological ‘need’ for drug. In this context, Ex-4 might be reducing the release of orexin associated with drug-craving, which in turn leads to an increase in OX1 expression in the NAcS. The orexin system is activated by stimuli of high motivational relevance such as stress, reward opportunities, or physiological need states [79]. Thus, treatment with an agent that can modulate the physiological state of the rat from ‘hungry’ (i.e. craving) to ‘sated’ (i.e., no motivation to seek), such as Ex-4, is effective in reducing relapse to opioid seeking in rats. As such, GLP-1 R agonists have great potential as a novel treatment for opioid misuse and addiction in humans, especially since these drugs already are FDA-approved for the treatment of type II diabetes mellitus and obesity in humans [40-42]. Indeed, given the urgency of the situation, the effort to test whether treatment with a GLP-1R agonist can reduce craving and relapse in humans with an opioid use disorder is under way.

## ACKNOWLEDGMENTS AND DISCLOSURES

The authors thank the National Institute on Drug Abuse for generously providing heroin. Support for this research was provided by NIH grants DA009815 (to PSG). Support also was provided by a grant from the Pennsylvania Department of Health, Tobacco Settlement Funds SAP# 410007972 (to PSG) and by the Elliot S. Vesell Professorship in Pharmacology (to KEV). Part of these data have been presented at the annual meeting of the Society for the Study of Ingestive Behavior (SSIB). We thank Sarah Ballard for her technical assistance.

This manuscript was submitted as preprint on bioRxiv server.

The authors report no financial interest or conflicts of interests.

