## Supplemental Material for "Glucagon-like peptide-1 receptor agonist, exendin-4, reduces reinstatement of heroin seeking behavior in rats"

### Subjects

The subjects were 55 outbred male Sprague-Dawley rats delivered from Charles River (Wilmington, MA) at approximately 90 days of age, weighing between 300 – 400g at the start of the experiment. All subjects were housed individually in standard, suspended, stainless steel cages. The environment in the animal colony room had controlled humidity and temperature (21 °C), with a 12/12 hour light/dark cycle, and lights on at 7:00 am. All experimental manipulations were conducted starting 2 h into the light phase of the cycle. Following one week of acclimation to their home cages, rats were habituated to experimenter handling by daily weighing. Food and water were available ad libitum, except where noted otherwise. All studies were approved by the Pennsylvania State University College of Medicine, Institutional Animal Care and Use Committee and performed in accordance with the National Institutes of Health specifications outlined in their Guide for the Care and Use of Laboratory Animals.

### Solutions

Sodium saccharin was obtained from Sigma, St. Louis, MO, prepared 24 h prior to testing, and presented at room temperature. The heroin dose was prepared by dissolving 0.075 g of heroin in 250 ml of 0.9% saline. A stock solution (4.8 µg/ml) of Exendin-4 (Tocris) was made by dissolving one milligram in 208 ml of 0.9% saline. Fifteen ml of the stock solution were diluted in 15 ml of 0.9% saline to make the 2.4 µg/ml solution. Stock solution, injection doses, and filled syringes were maintained at -18 °C and thawed before use. All light-sensitive solutions were covered in aluminum foil.

### Self-Administration Catheter

### Jugular Catheterization Surgery

Rats were anesthetized with intramuscular (im) ketamine (70 mg/kg) and xylazine (14 mg/kg) and then implanted with custom-made intravenous jugular catheters as described previously [1]. Following surgery, rats received two days of post-op care consisting of a daily sc injection of the NSAID, carpofen, and a full week to recover.

Carprofen treatment was continued when indicated. Maintenance of catheter patency included flushing of catheters using heparinized saline (0.2 ml of 30 IU/ml heparin) daily. Catheter patency was verified when necessary using 0.3 ml of propofol (Diprivan 1%). All studies were approved by the Pennsylvania State University College of Medicine Institutional Animal Care and Use Committee and performed in accordance with the National Institutes of Health specifications outlined in their Guide for the Care and Use of Laboratory Animals.

**Apparatus**

Testing was conducted in 34 self-administration chambers as previously described by Grigson and Twining [1]. Each chamber measured 30.5 cm in length, 24.0 cm in width, 29.0 cm in height, and was equipped with three retractable sipper tubes that entered the chamber through three holes. A stimulus light was located above each hole. A lickometer circuit was used to monitor licking on all three spouts. In addition, each chamber was equipped with a house light (25 W), a speaker for white noise and a tone generator. Collection of data and events in the chamber were controlled on-line with a Pentium computer using programs written in the Medstate notation language (MED Associates, Inc., St. Albans, VT).

**Molecular analysis**

Gene expression analysis

NAc shell was homogenized and RNA was isolated using the Allprep DNA/RNA kit (Qiagen). Complimentary DNA (cDNA) was generated from the isolated RNA using Random hexamer primers (Invitrogen) and Superscript III Reverse Transcriptase (Invitrogen). Messenger RNA expression was analyzed using real time quantitative polymerase chain reaction (RT-qPCR) Quantstudio 102k flex. The $2^{-\Delta\Delta CT}$ method was employed to assess relative gene quantification with beta actin as the endogenous control [2] and a set of naïve rats without behavioral manipulation was used to normalize gene expression. Quantitative PCR analysis of targets of interest was performed using standard laboratory methods, with 384-well optical plates, TaqMan

Assay-On-Demand (Applied Biosystems, Foster City, CA, USA) gene-specific primers/probe assays and a 7900HT Sequence Detection System (Applied Biosystems). Gene expression assays included Orexin Receptor 1 (Hcrtr1, Rn0056995_m1), D2 (d2r, Rn00561126_m1), Leptin receptor (Lepr, Rn00565158_m1) and GLP-1 receptor (glp-1r, Rn00562406_m).

**Data analysis**

Subjects that lost catheter patency were removed from the subsequent phase of the experiment. All data were analyzed using Prism version 5.00 for Windows, GraphPad Software (La Jolla California USA). Mixed factorial Analysis of Variance (ANOVAs), followed by Newman-Keuls post hoc tests, were used to compare different groups. In addition, Student-t tests were used when comparing only two means, with alpha set at 0.05.

Supplementary Results

**Body weight across abstinence days**

Body weights are shown in Figure S1 from the start of the experiment (time 0) through abstinence days 1–16. Results show that, while heroin self-administration was associated with lower body weight overall, it was not significantly impacted by daily treatment with Ex-4 throughout the abstinence period. Support for these conclusions was provided by conducting a 2 x 2 x 16 mixed factorial ANOVA varying drug (saline or heroin), treatment (vehicle or Ex-4), and abstinence days 1–16. Results revealed a significant main effect of drug ($F_{1,47}$=22.96, p<0.0001), with rats that self-administered heroin weighing significantly less than rats that self-administered saline. Additionally, while there was a trend for Ex-4-treated rats to weigh less, neither the main effect of treatment ($F_{1,47}$=1.47, p=0.23), the group x treatment (F<1.0), or the group x treatment x day (F< 1.0) interactions were statistically significant. Hence, treatment with the present dose of Ex-4, then, did not have an effect on body weight in either the saccharin-heroin group or in the saccharin-saline controls.

**Figure S1**

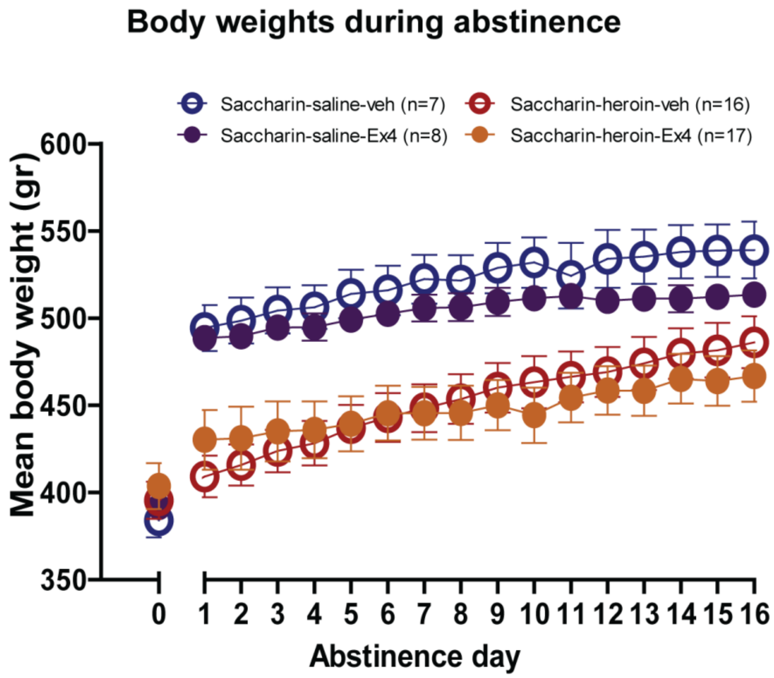


**Figure S1. Body weight across experiment.** Mean± SEM body weight in grams for saccharin-saline rats treated with Veh (n=7) or Ex-4 (n=8) and saccharin-heroin rats treated with Veh (n=16) or Ex-4 (n=17). Day 0 represents average body weight at the beginning of acquisition and days 1 to 16 represent abstinence days.
